# Bimodal fibrosis in a novel mouse model of bleomycin-induced usual interstitial pneumonia

**DOI:** 10.1101/2021.03.18.435059

**Authors:** Yoko Miura, Maggie Lam, Jane E. Bourke, Satoshi Kanazawa

**Affiliations:** Department of Neurodevelopmental Disorder Genetics, Nagoya City University Graduate School of Medical Sciences, Nagoya, Aichi, Japan; Department of Pharmacology, Biomedicine Discovery Institute, Monash University, Clayton, Australia

**Keywords:** bleomycin, honeycombing, idiopathic pulmonary fibrosis, interstitial pneumonia, usual interstitial pneumonia, fibrosis

## Abstract

Idiopathic pulmonary fibrosis (IPF) is pathologically classified by usual interstitial pneumonia (UIP). Conventional bleomycin models used to study pathogenic mechanisms of pulmonary fibrosis display transient inflammation and fibrosis so their relevance to UIP is limited. We developed a novel chronic <u>i</u>nduced-UIP (iUIP) model, inducing fibrosis in D1CC×D1BC transgenic mice by intra-tracheal instillation of <u>b</u>leomycin mixed with <u>m</u>icrobubbles followed by <u>s</u>onoporation (BMS). A bimodal fibrotic lung disease was observed over 14 weeks, with an acute phase similar to nonspecific interstitial pneumonia (NSIP), followed by partial remission and a chronic fibrotic phase with honeycombing similar to UIP. In this secondary phase, we observed poor vascularization despite elevated PDGFRβ expression. γ2PF- and MMP7-positive epithelial cells, consistent with an invasive phenotype, were predominantly adjacent to fibrotic areas. Most invasive cells were Scgb1a1 and/or keratin 5 positive. This iUIP mouse model displays key features of IPF and has identified potential mechanisms contributing to the onset of NSIP and progression to UIP. The model will provide a useful tool for the assessment of therapeutic interventions to oppose acute and chronic fibrosis.

**Summary blurb:** A novel mouse model of induced-usual interstitial pneumonia shows bimodal fibrosis with honeycombing, similar to chronic lung disease seen with idiopathic pulmonary fibrosis.

## INTRODUCTION

Interstitial pneumonia (IP) is a lung dysfunction with expanding fibrotic lesions due to the accumulation of extracellular matrix (ECM). Idiopathic pulmonary fibrosis (IPF) is classified as one of the most serious chronic IPs leading to the loss of pulmonary function (Antoniou et al 2014). The histopathology of IPF is defined by usual interstitial pneumonia (UIP), which is derived from various features such as honeycombing and bronchiolization of alveoli together with severe fibrosis (Hashisako & Fukuoka 2015, Raghu et al 2018, Smith et al 2013). IPF is also associated with the loss of endothelial cells and microvasculature causing pulmonary hypertension (Farkas et al 2009, Nathan et al 2007). The underlying mechanisms of these pathological changes in IPF are not yet fully defined, findings from studies of both clinical samples and short-term bleomycin models have implicated repetitive trauma of epithelial cells and dysregulation of repair processes mediated by such as transforming growth factor-β (TGF-β) in disease progression (Hashisako & Fukuoka 2015).

The development of pulmonary fibrosis is most commonly studied in animal models using bleomycin, an oligopeptide antibiotic produced by *Streptomyces verticillus* (Hay et al 1991, Organ et al 2015). Although bleomycin has established chemotherapeutic effects against cancer, its side effects in patients include pathologic changes consistent with IPF (Adamson 1976). In animal models, administration of single or multiple doses of bleomycin by intratracheal (i.t.) instillation, osmotic pump, intravenous route, or intranasal delivery induces pulmonary fibrosis, with significant dose-dependent mortality (B. et al 2013, Jenkins et al 2017, Mouratis & Aidinis 2011). Bleomycin mediates DNA double-strand breaks (DSBs) and thus leads to the induction of lung injury with severe inflammation (Mouratis & Aidinis 2011). The half-life of bleomycin is from a few hours up to 21 hours in the body. It triggers events seen in the onset of pulmonary fibrosis, namely epithelial injury and alveolar infiltration of lymphoid cells such macrophages (Cooper 2000, Dorr 1992). In most animal studies, bleomycin-treated mice are analyzed 14 to 28 days after a single intra-tracheal dose. Acute inflammatory changes in C57BL/6 mice over this relatively short-time frame resemble those seen in lung diseases like nonspecific interstitial pneumonia (NSIP), but usually occur without the development of chronic symptoms unless repeated doses of bleomycin are administered (Satoh et al 2017). Assemble of potential therapeutic agents are usually assessed in these short-term bleomycin models without monitoring IP biomarkers such as surfactant protein-D (SP-D) or micro-CT (Murata et al 2010). As such, it is often difficult to distinguish between the effects of treatment and spontaneous recovery (Moeller et al 2008). Of note, current models of single bleomycin instillation in C57BL/6 mice do not generally result in the long-term histological characteristics seen with UIP in patients with IPF except recently published data by Redente *et al* (B. et al 2013, Borzone et al 2001, Limjunyawong et al 2014, Redente et al 2021, Redente et al 2011, Tashiro et al 2017).

Although the cause of IPF remains unknown, findings from studies of both clinical samples and short-term bleomycin models have implicated repetitive trauma of epithelial cells and dysregulation of repair processes mediated by transforming growth factor-β (TGF-β) in disease progression (Hashisako & Fukuoka 2015, Jiang et al 2020). Of particular note, blockage of TGF-β signaling resulted in attenuation of fibrotic responses during wound healing in bleomycin-induced pulmonary fibrosis (Degryse et al 2011), informing the current first-line treatment of IPF with pirfenidone and nintedanib (Flaherty et al 2019, King et al 2014).

To address these limitations, we have developed a new model of UIP, using D1CC^+/+^×D1BC^+/+^ transgenic mice (hereafter D1CC×D1BC tg), bred on a DBA/1J background. These mice were bred by crossing IP-susceptible D1CC and D1BC tg mice, which express class II transactivator (CIITA as an MHC class II transcriptional co-activator) and B7.1 (co-stimulatory signaling molecule), respectively (Fontes et al 1999, Kanazawa et al 2000, Miura et al 2019). D1CC, D1BC, and D1CC×D1BC tg mice have chronic joint inflammation, as seen in patients with rheumatoid arthritis (RA) with additional interstitial lung disease (ILD) (Kanazawa et al 2006, Miura & Kanazawa 2020, Miura et al 2019, Terasaki et al 2019). After induction of RA by low-dose type II collagen, DBA/1J mice did not develop pulmonary inflammation, while the tg mice had increased serum SP-D levels and chronic deposition of ECM in the lung with higher susceptibility compared to C5BL/6 mice (Schurgers et al 2012, Terasaki et al 2019). Histopathology of lung tissues in mice with joint inflammation was characterized as NSIP, defining this as a model of RA-ILD.

Given this background, we hypothesized that bleomycin would induce features of NSIP and potentially UIP in D1CC×D1BC mice. To facilitate the development of a chronic model, we utilized a novel method to deliver a lower single dose of <u>b</u>leomycin than usually administered mixed with <u>m</u>icrobubbles and <u>s</u>onoporation (hereafter called BMS) to minimize mortality while increasing drug delivery to the lung (Okada et al 2005, Tashiro et al 2017, Xenariou et al 2010, Yamaguchi et al 2017). This BMS mouse model induced bimodal fibrosis, alveolar bronchiolization and altered vascularity, mimicking key features of UIP in IPF, and enabled the characterization of specific phenotypes of the resident cells contributing to these pathological changes.

## METHODS

### Mice and BMS administration

D1CC×D1BC tg mice, bred on a DBA/1J background, were housed in a pathogen-free animal care facility of Nagoya City University Medical School in accordance with institutional guidelines (Kanazawa et al 2006, Miura et al 2019). Mice were anesthetized with isoflurane and the chest hair was shaved for sonoporation. Bleomycin (0.512 mg/mL in normal saline, Nippon Kayaku) was mixed with an equal amount of microbubbles (Ultrasound Contrast Agent SV-25, NepaGene) and administered via the intra-tracheal (i.t.) route (40 μl/mouse, 1.28 mg/kg) prior to sonoporation on the chest by 1.0 W/cm^2^ for 1 minute (Sonitron GTS Sonoporation System, NepaGene). Induction of IP was monitored by measurement of serum SP-D levels.

The selected bleomycin dose was the mid-range of dose-ranging pilot studies using 0.96-1.6 mg/kg, also given in microbubbles with sonoporation. These BMS doses resulted in SP-D levels of >1000 ng/ml with low mortality, while lower BMS doses or microbubbles or sonoporation alone did not induce IP (Fig S1).

### ELISA

Serum was collected from the external jugular vein of each mouse from week 0 to 14 after BMS administration, aliquoted and stored at −80°C. Quantitation of serum SP-D concentrations was determined using ELISA according to the manufacturer’s instructions (Rat/Mouse SP-D kit, Yamasa).

### Western Blot

Lung samples were homogenized in RIPA buffer, containing 20 mM Tris-HCl, pH 7.4, 150 mM NaCl, 1% Triton, 0.5% deoxycholate sodium, 0.1% sodium dodecyl sulfate, 1 mM EDTA, 10 mM BGP, 10 mM NaF and protease inhibitor cocktail. The extracts were sonicated for 10 minutes, and centrifuged at 12,000 rpm for 15 minutes. The supernatants were separated on 8, 14, or 18% SDS-PAGE, and transferred onto polyvinylidene difluoride membranes. Protein-transferred membranes were blocked in 5% nonfat milk/ TBS-T buffer at room temperature for 60 minutes. For Western blot, the following primary antibodies were used: rabbit anti-collagen (Boster biological technology), rabbit anti-αSMA, rabbit anti-vimentin, rabbit anti-PDGFRα, rabbit anti-PDGFRβ and rabbit anti-E-cadherin (Cell Signaling Technology), goat anti-SP-D, mouse anti-TGF-β and goat anti-Serpin F1/PEDF (R&D Systems), rabbit anti-Uteroglobin, Rabbit anti-CD31 and rabbit anti-VEGFA (Abcam), rabbit anti-SP-A, mouse anti-PDGFA and mouse anti-Tubulin-α (Santa Cruz), rabbit anti-SP-C (Hycult Biotech) and rabbit anti-PDGFB (Bioss antibodies). ECL™ anti-rabbit or anti-rat IgG, horseradish peroxidase linked antibodies were used as a secondary antibody (GE Healthcare). Each signal was detected using Immunostar Zeta or LD (Fuji film) and the Amersham Imager 600 series (GE Healthcare). Statistical analysis of the expression levels of each protein were evaluated by ImageJ, Fiji.

### Analysis of lung sections for cellular and pathological markers

Lungs were collected at 0, 6, 12, 24h, days 3, 7, and weeks 2, 4, 6, 8, 10, 14 weeks after BMS administration, fixed overnight in 4% paraformaldehyde diluted in PBS then embedded in paraffin before 2 μm thick sections were cut. De-paraffinized sections were stained with hematoxylin-eosin (H&E) or Masson’s trichrome to assess inflammation and fibrosis, or subjected to conventional TUNEL assay according to the manufacturer’s instructions to detect apoptotic cells (Mebstain Apoptosis Tunel kit, MBL). For immunohistochemistry, the de-paraffinized sections were stained with the following primary antibodies: rabbit anti-E-cadherin, rabbit PDGFRα, rabbit PDGFRβ, rabbit anti-αSMA and rabbit anti-MMP7 (Cell Signaling Technology), mouse anti-phospho-H2AX and rabbit anti-S100A4 (Merck Millipore), rabbit anti-CD31 and rabbit anti-VEGFA (Abcam), rabbit anti-SP-C (Hycult Biotech), rat anti-Podoplanin (MBL), rat anti-F4/80 (Bio-Rad), rabbit anti-CD3 (Genemed Biotechnologies), rat anti-B220/CD45R (Aviva Systems Biology), rabbit anti-collagen I (Novus Biologicals), rat anti-Ki67 (Dako), mouse anti-γ2pf (Funakoshi). Histofine Simple Stain Mouse MAX-PO secondary antibodies were used and visualized using the Histofine SAB-PO (M) kit (Nichirei Biosciences). For immunofluorescence, the Opal 4-color fluorescent IHC kit was used (PerkinElmer). All images were captured and assessed for statistical analyzes using BZ-X analyzer (Keyence) and ImageJ, Fiji (Schindelin et al 2012).

### Quantitative analysis of phospho-H2AX (γH2AX)-positive cells

Immunohistochemical staining of γH2AX was performed to assess the percentage of DNA-damaged cells at day 0, 3, 7 and 14 after BMS administration, using Hematoxylin as a counter stain. Five images (a magnification of ×200) from each lung section were captured randomly and the percentage of γH2AX-positive cells was calculated by ImageJ, Fiji. For calculation of the percentage of each lung cell type within the γH2AX-positive cells, multi-color immunohistochemistry was performed for γH2AX, podoplanin (AEC1), SP-C (AEC2), E-cadherin (bronchial epithelial cell), S100A4 (fibroblast). Ten images (at a magnification of ×200) from each lung section were captured randomly and analyzed by hybrid cell count (Keyence).

### Ashcroft Score

The severity of lung fibrosis and destruction of lung structure including formation of honeycomb structure was evaluated by Ashcroft score using scanned Masson’s trichrome stained sections (Ashcroft et al 1988). For analysis, ten images from ×100 micrograph of whole lung sections were randomly selected from three mice for each day and week.

These randomly selected images were individually assessed for IP severity by M.Y. and S.K. in a blinded manner with an index of 0 to 8: 0 - Normal lung; 1 - Minimal fibrous thickening of alveolar or bronchiolar walls; 2-3 - Moderate thickening of walls without obvious damage to lung architecture; 4-5 - Increased fibrosis with definite damage to lung structure and formation of fibrous bands or small fibrous masses; 6-7 - Severe distortion of structure and large fibrous areas, evidence of “honeycomb lung”; 8 - Total fibrous obliteration of the field (Hubner et al 2008).

### Fibrosis ratio

Images showing the area of fibrosis represented in blue (ECM-deposition) by Masson’s trichrome staining were captured by BZ-X analyzer (Keyence) and analyzed by ImageJ, Fiji. Data was calculated as fibrosis ratio, with ECM area divided by total lung area.

### Histoindex

Multiple outcomes were measured in mice up to 14 weeks after BMS. A novel measure of lung fibrosis used the Genesis 200 (Histoindex, located in Pharmacology, Monash University) to visualize unstained lung sections using Second Harmonic Generation (SHG) to detect collagen as green light emission, with Two-Photon Excitation (TPE) Fluorescence-based microscopy showing lung tissue in red. Images were captured with laser settings adjusted to 0.65 at 20× magnification with 512×512 pixels resolution, then assessed by the Fibroindex application for morphological analysis of interstitial collagen area and fiber density within the tissue (Goh et al 2019).

### *In situ* hybridization

*In situ* hybridization for *Scgb1a1, Sftpc, Krt5, Col1a1*, and *Tgf-β* was performed using the RNAscope Multiplex Fluorescent Reagent Kit v2 (Advanced Cell Diagnostics), according to the manufacturer’s instructions.

### Soluble collagen assay

Soluble collagen content from whole lung extracts was determined by the Sircol assay (Biocolor), according to the manufacturer’s instructions. Sircol dye bound to collagen was evaluated by a microplate reader at 555 nm.

### Micro-computed tomography

Mice were anesthetized with isoflurane and the abdominal and chest cavities were opened. The lung vasculature was perfused with PBS via the right ventricle, then fixed using 4% paraformaldehyde in PBS. Microfil^®^(Flow Tech Inc.) was then injected via the same route and incubated for 1h at room temperature (Phillips et al 2017). Whole lung was harvested and fixed again overnight in 4% paraformaldehyde in PBS. All lungs were stored in 70% ethanol at 4°C until micro-CT scanning. All micro-CT images were acquired with a SkyScan 1276 (Bruker). Five images for each lung were randomly selected adjacent to the subpleural area and the density of blood vessels was calculated.

### Statistical analyzes

The mortality rate in the iUIP mouse model was calculated by the Kaplan-Meier method (Prism9, GraphPad). Differences between non-instillation (0w) and the other groups were evaluated by one-way analysis of variance (ANOVA) followed by Dunnett’s test for parametric data and Dunn’s test for nonparametric data. Tukey’s post hoc test was used for multiple comparisons of the percentage of γH2AX-positive cells in different cell types. Values of *p* < 0.05 were considered statistically significant.

## RESULTS

### BMS induces chronic IP

Following a single i.t. instillation of BMS in D1CC×D1BC mouse (Fig 1A), disease progression was monitored by the IP-biomarker, serum SP-D (Miura et al 2019). The peak SP-D concentration after BMS was > 2,500 ng/ml at week 2, approximately three-fold higher than after i.t. instillation of the same dose of bleomycin alone (Fig 1B). Serum SP-D levels declined markedly in both groups by 6 weeks, with only the BMS group still slightly elevated above the cut-off value (>53.9 ng/ml) after 8 weeks (Figure 1B inset). Sonoporation either alone or in combination with microbubbles, did not induce sufficient trauma to cause the development of IP, as evidenced by its lack of effect on serum SP-D, while lower doses of BMS were also ineffective in the tg mice (Fig S1). Total lung expression levels of SP-D were decreased when serum SP-D was at its peak during disease progression. (Fig 1C). The patterns of serum and lung SP-D levels suggest that the maximal leakage from pulmonary parenchyma into the blood coincides with acute BMS-induced inflammation. Thus, there was incompatibility between serum SP-D and total SP-D in the whole lung during disease progression. We also examined total lung expression levels of the other C-type lectins, SP-A and -C by Western blot (WB). SP-A levels were lower at week 2 compared to untreated controls, and tended to recover to at or above control levels at week 14 in the chronic phase, while SP-C levels were slightly increased (Figs 1D, 1E). Finally, the mortality with this relatively low bleomycin dose was less than 20%, with only 2 deaths in a total of 14 mice over 14 weeks (Fig 1F).

**Figure 1.**
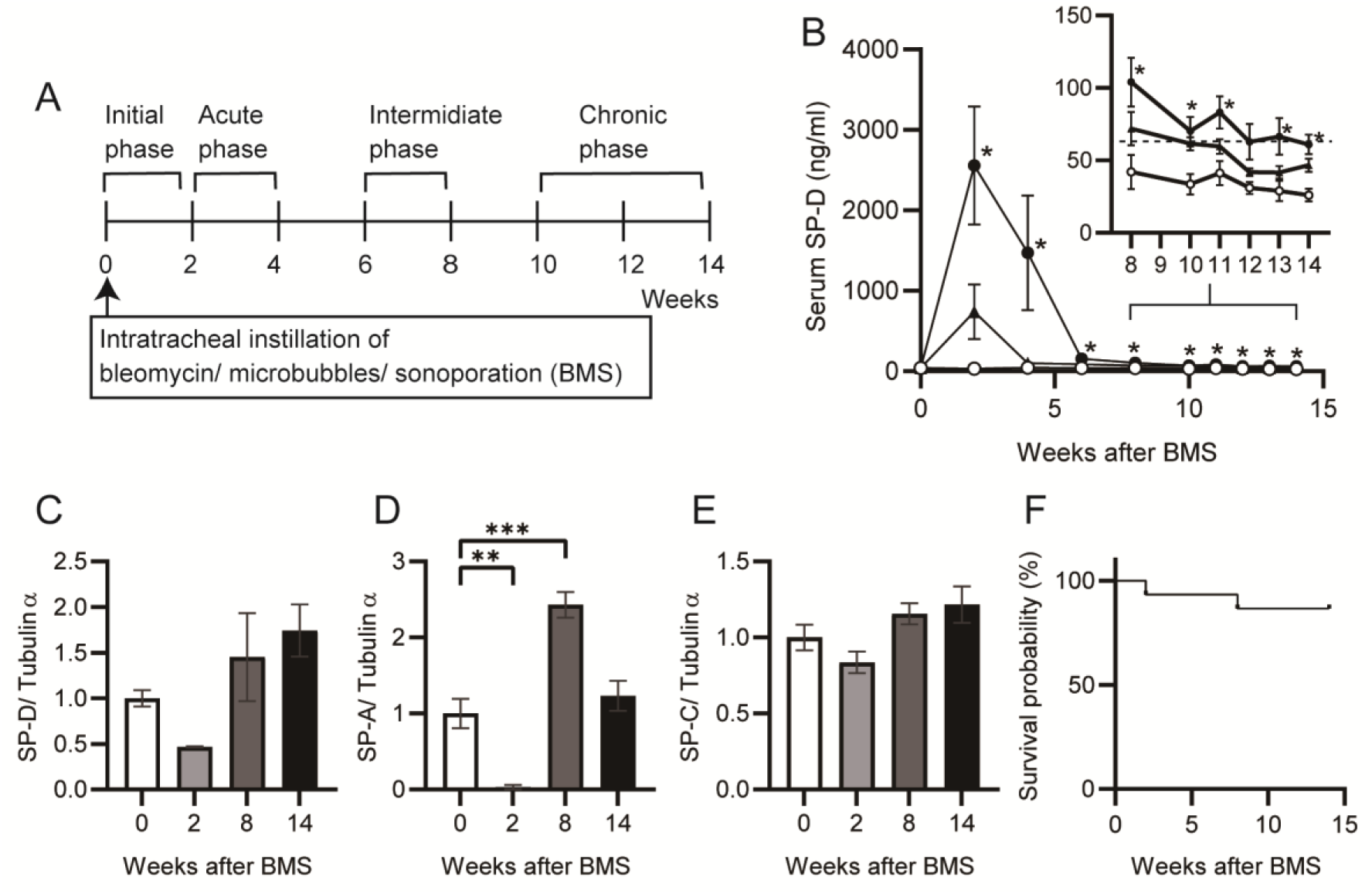
Serum and lung SP-D levels during BMS-induced disease progression. (A) Timeline following i.t. instillation of bleomycin (1.28 mg/kg) with microbubbles followed by sonoporation (BMS) in D1CC×D1BC mice, showing initiation, acute, intermediate, and chronic phases of IP progression. (B) Serum SP-D levels over 14 weeks following BMS (closed circle), bleomycin alone (closed triangle), or vehicle (open circle) in D1CC×D1BC mice evaluated by ELISA. The dotted line in inset graph indicates serum SP-D cut-off value (53.9 ng/ml). Data are mean ± S.E. of seven mice at each timepoint. Asterisks show *\*P* < 0.05, compared with vehicle. (C-E) SP-D (C), SP-A (D), and SP-C (E) in whole lung extracts at weeks 0, 2, 8, and 14 measured by WB. The relative signal intensity from densitometrical analysis was calculated using ImageJ Fiji, with tubulin-α used as a loading control. Data are mean ± S.E. from three mice per group. Asterisks show \*\**P* < 0.01, and \*\*\**P* < 0.001, compared with 0 weeks. (F) Kaplan-Meier analysis of mouse survival post-BMS from weeks 0 to 14 of 14 mice.

### BMS induces DNA damage lung restricted to AEC1, AEC2 and bronchiolar epithelial cells

We examined whether bleomycin induced cell death during disease progression. Apoptotic or necrotic cells were observed in lung alveoli at 6 to 24h (Fig 2A). Consistent with bleomycin-induced cell death occurring mainly within one day of administration, cell death did not increase after day 3 when the percentage of dead cells was less than 0.5% of total lung cells (Fig S2) (Hagimoto et al 1997). The local effects of BMS seen on lung epithelial cells were not evident either in lung stromal cells or other tissues such as skin and liver (Fig S3).

**Figure 2.**
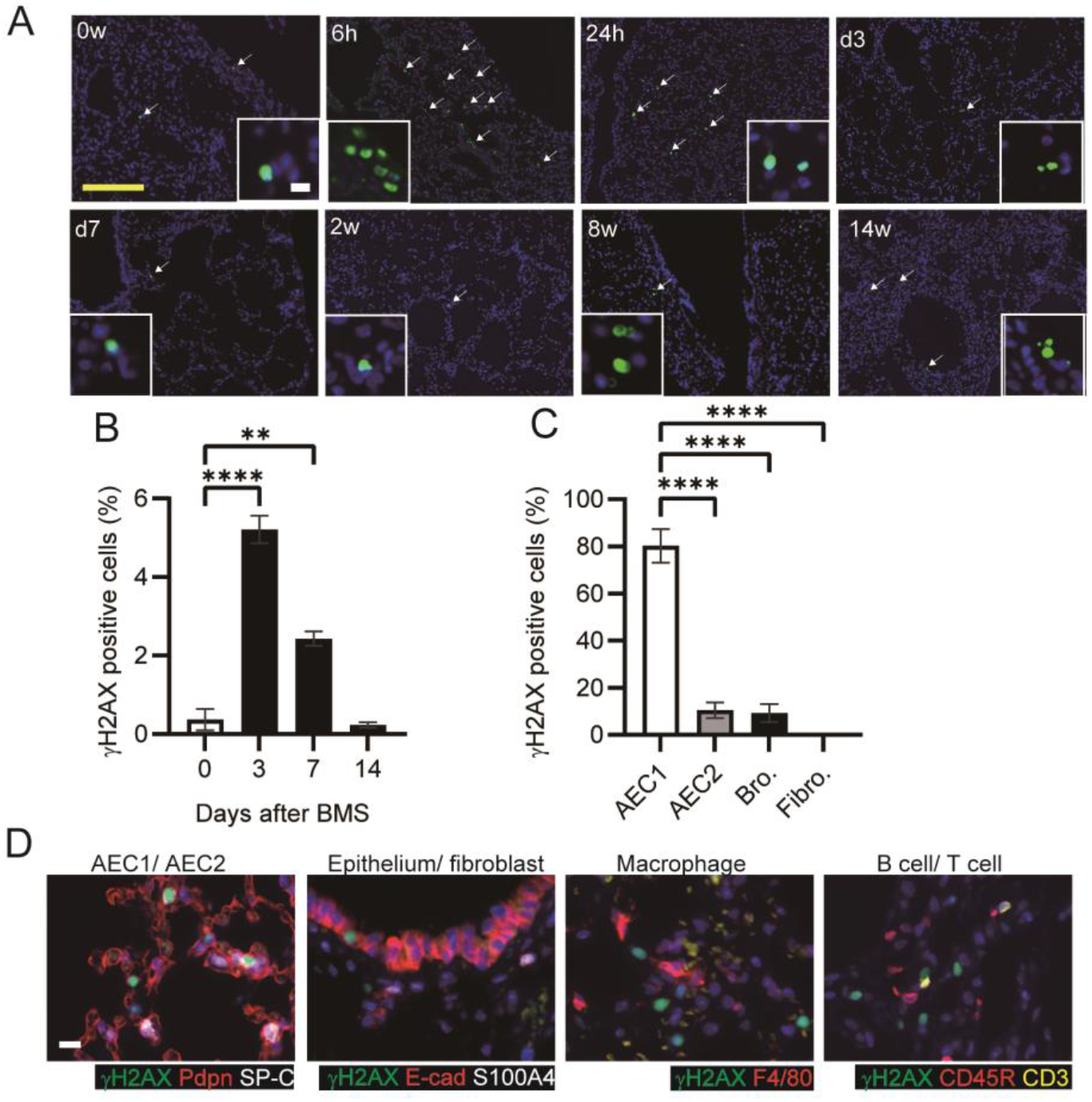
DNA damage in AEC1, AEC2 and bronchioles, but not in fibroblasts. (A) TUNEL assays in mouse lungs at 0, 6, 24h, days 3, 7, and weeks 2, 8, 14 after BMS administration. Arrows indicate cell death. (B) Percentage of γH2AX-positive cells at days 0, 3, 7 and 14 in mice treated with BMS. Images were captured and analyzed from five randomly selected fields on each slide at a magnification of 200×. Data are mean ± S.E. from five mice per group. Asterisks show *\*\*P*< 0.01, *\*\*\*\*P*< 0.0001, compared with 0 weeks. (C) Percentage of γH2AX-positive cells in AEC1 (podoplanin), AEC2 (SP-C), bronchiolar epithelium (E-cadherin, Ecad), and fibroblasts (S100A4) at day 3. Images were captured and analyzed from ten randomly selected fields on each slide at a magnification of 200×. Data are mean ± S.E. from three mice per group. Asterisks show \*\*\*\**P* < 0.0001, compared with the percentage of γH2AX-positive cell in different cell type. (D) Immunohistochemical staining of γH2AX (green), podoplanin (Pdpn, red), SP-C (white), E-cadherin (E-cad, red), S100A4 (white), F4/80 (red), CD45R (red), and CD3 (yellow). Scale bar = 100 μm (yellow) and 10 μm (white).

Bleomycin is a radical transfer agent, which induces single- or double-strand DNA breaks in the nucleus. We examined the percentage of γH2AX-positive cells as a marker of this DNA damage at days 3, 7 and 14. Approximately 5% of total lung cells were γH2AX positive at day 3, with numbers decreasing to similar levels as in untreated controls by week 2 (Fig 2B). Thus, bleomycin-induced DNA breaks may be limited to the first few days. Next, we investigated which cell types were susceptible to bleomycin. The vast majority of γH2AX positive cells were podoplanin-positive AEC1s at day 3 (Fig 2C). The bronchiolar epithelium and AEC2 also displayed significant susceptibility to bleomycin, but fibroblasts and lymphocytes were rarely γH2AX positive (Fig 2D). Since 90 to 95% of the lung surface is covered with AEC1s, the bronchiolar epithelium and AEC2 may be relatively more sensitive to bleomycin-induced DNA breaks than AEC1 while fibroblasts may be resistant (McElroy & Kasper 2004).

### BMS induces bimodal fibrosis characterized by acute inflammation and chronic honeycombing

To measure fibrosis in BMS-induced D1CC×D1BC mice, we assessed Masson’s Trichrome stained sections at selected time-points over 14 weeks (Figs 3A and S4). Analysis of digital images by ImageJ, Fiji was performed to obtain fibrosis area ratio values (Fig 3B). Pulmonary fibrosis was evident by week 2, with peaks observed at week 2 to 6 in the acute phase and also at week 14 in the chronic phase. Some level of fibrosis was retained during remission with secondary diffuse fibrosis observed at the pleural surface at week 8 (Fig S4). Notably, severe fibrosis with honeycomb structure, but not marked infiltration of lymphoid cells was seen in the chronic phase. This is because the percentage of macrophages and T cells increased at week 2, but infiltrated cells were restored to control levels at weeks 8 and 14 (Fig S5).

**Figure 3.**
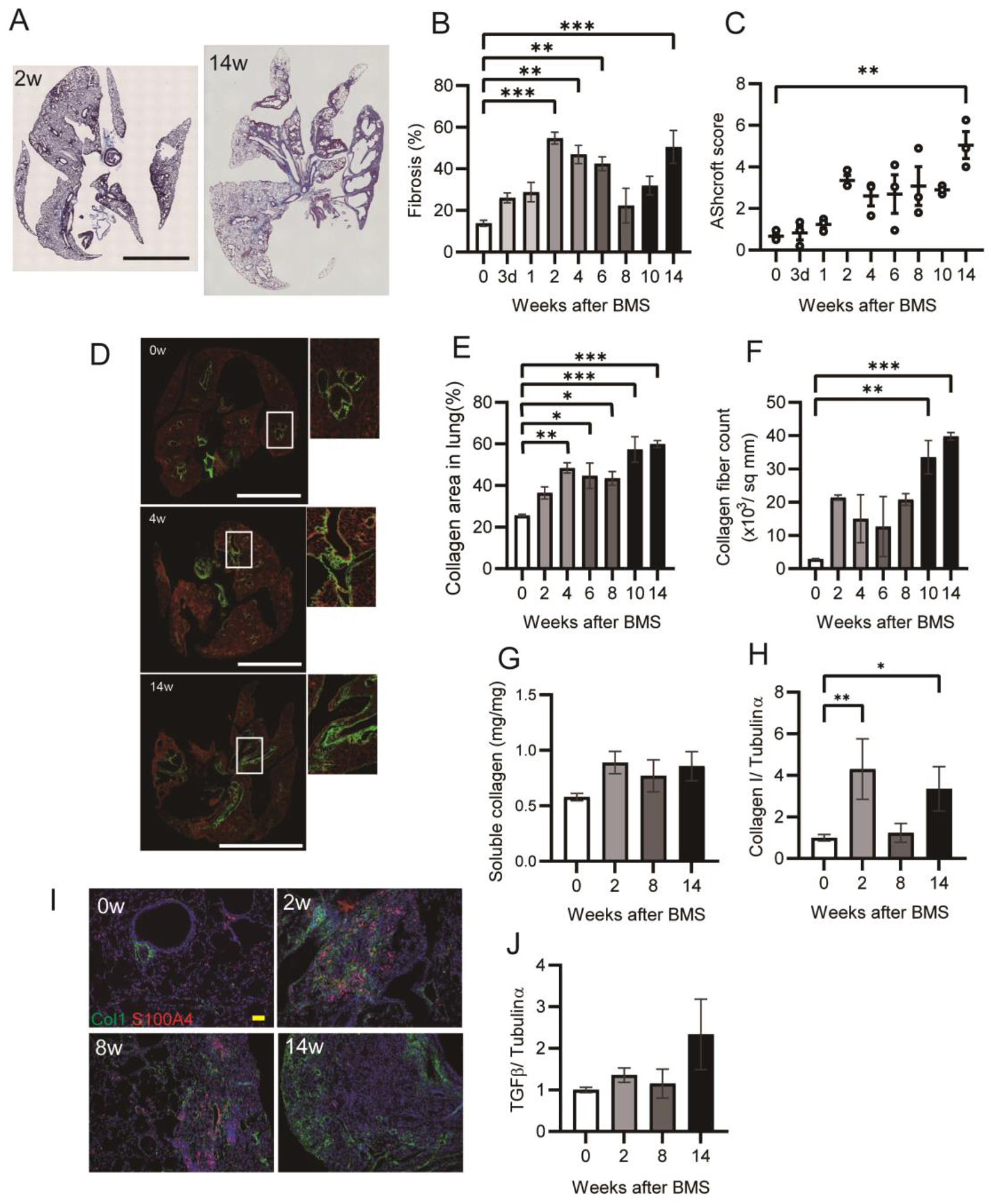
Bimodal fibrotic stages in IP progression. (A) Representative histopathological macro- and micro-graphs of Masson’s trichrome stained sections from BMS-induced D1CC×D1BC mice at weeks 2 and 14. (B) Quantitative analysis of fibrosis from Masson’s trichrome stained sections as ratios of ECM area (blue) to total lung area, as calculated by ImageJ, Fiji. n = 6-7 per each group. (C) Quantitative analysis of Ashcroft score (averaged from scores by two individuals, YM and SK, performed in a blinded manner). n = 6-7 per group. (D) Representative images at weeks 0, 4 and 14 captured with Histoindex show the fibrotic area (mostly collagen I and III, green) and the entire lung area (cell structure, red). (E) Collagen deposition and (F) collagen fiber density in 2 μm thick unstained specimens captured by Histoindex and analyzed using Fibroindex software. n =3 per each group. (G) Quantitative analysis of total soluble collagen using sirius red. n = 5 per each group. (H) Col1 expression determined by WB and ImageJ, Fiji. n = 3 per each group. (I) Immunohistochemical staining of Col1 (green) and S100A4 (red) at weeks 0, 2, 8, and 14. (J) TGF-β expression determined by WB and ImageJ, Fiji. Scale bars in (A) = 5 mm (black) and (F) =100 μm (white) and (I) = 50 μm (yellow). (G, H and J) n = 3 per each group. Data are presented means ± S.E. Asterisks show \**P* < 0.05, \*\**P* < 0.01, and \*\*\**P* < 0.001, compared with week 0.

The highest Ashcroft score for lung fibrosis was associated with the honeycomb structure seen in the specimens at week 14 (Fig 3C). A similar bimodal pattern of fibrosis was observed using Histoindex stain-free imaging to detect collagen combined with Fibroindex analysis to determine collagen area and collagen fiber density (Figs 3D to 3F).

The total amount of fibrillar collagen, measured by a Sircol assay for soluble collagen, was also increased at weeks 2, 8 and 14 (Fig 3G). By comparison, only weak IP was also observed in D1CC alone, D1BC alone, and their genetic background DBA/1J mice (all scored <3 at 14 weeks), suggesting minimal susceptibility for fibrosis in response to BMS administration in DBA/1J mouse (Fig S6)

Expression of collagen type I (Col1), measured by WB, showed a similar pattern to measures of fibrosis (Fig 3H). Immunohistochemical staining and WB confirmed that the increased αSMA was localized to fibrotic areas evident after BMS administration (Fig S7). We performed double-immunohistochemical staining for Col1 and S100A4 (also called fibroblast specific protein 1 (FSP1)). The expression of both proteins overlapped at week 2 and 8 in the fibrotic area, however less S100A4 expression was observed in the Col1-rich fibrotic area at week 14 (Fig 3I). Expression of TGF-β, a robust stimulator of fibrosis that also inhibits cell proliferation, was measured in the whole lung by WB. This was elevated only at week 14, but not at weeks 2 and 8 (Fig 3J). Overall, our analysis established a highly consistent pattern of bimodal fibrosis in the BMS model.

### Extension of fibroblasts in fibrotic foci in chronic phase

Further characterization of fibrotic changes in BMS-induced D1CC×D1BC mice was performed by WB of whole lung. Vimentin (fibroblast marker) and PDGFRβ (pericytes) were significantly increased at week 2, and decreased at later timepoints, while expression of PDGFRα (fibroblasts) only showed a gradual decline (Figs 4A to 4C). Next, we analyzed the expression of PDGFR ligands, PDGFα and PDGFβ, in the whole lung. Only PDGFβ was elevated at week 2, but this was not maintained during disease progression (Figs 4D, 4E). This suggests that PDGFβ-mediated growth of fibroblasts may be limited to the acute phase. To further focus on fibrotic foci at week 14, we performed multiple immunohistochemical analyses using antibodies against E-cadherin (epithelial cell marker), SP-C (epithelial cells and AEC2s) and Col1 (fibroblasts). A Col1^+^ fibrotic area was localized adjacent to, but distinct from, another area consisting of E-cadherin+/ SP-C^+^ epithelial cells (Fig 4F). Further analysis in this Col1-positive region showed that PDGFRα and PDGFRβ double-positive rather than PDGFRα single-positive areas were predominant (Fig 4G). A limited number of PDGFR-positive fibroblasts expressed Ki-67 simultaneously (Fig 4H). These data suggest that in the absence of increased PDGF at week 14, most fibroblasts tend to aggregate or extend rather than continue to proliferate in the fibrotic region.

**Figure 4.**
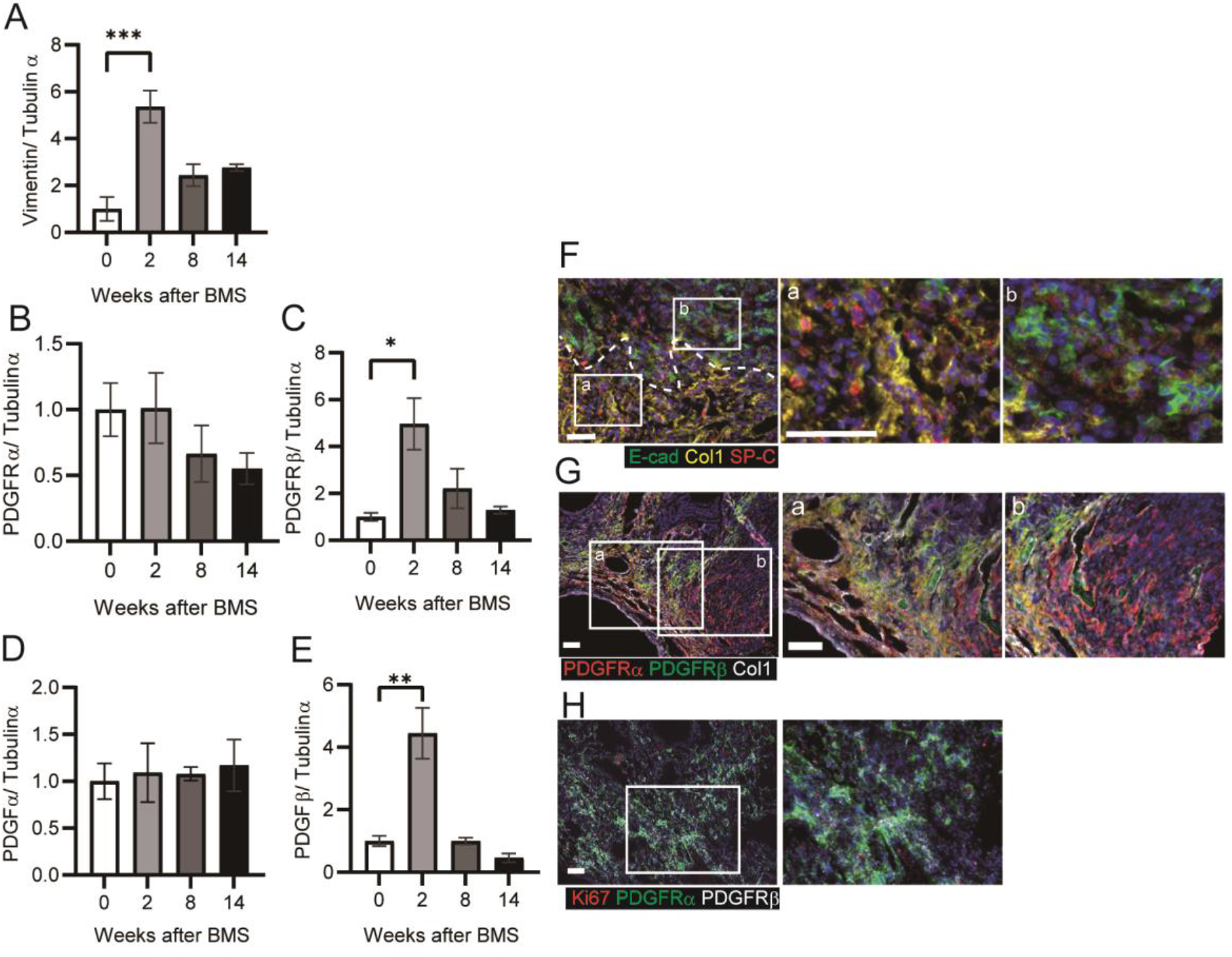
Extension of fibroblasts rather than proliferation in fibrotic foci in chronic phase. (A-E) Expression of vimentin (A), PDGFRα (B), PDGFRβ (C), PDGFα (D), and PDGFβ (E) at weeks 0, 2, 8, and 14 determined by WB and ImageJ, Fiji. Data are presented means ± S.E. of three mice for each week. Asterisks shows \**P* < 0.05, \*\**P* < 0.01, and \*\*\**P* < 0.001, compared with 0 weeks. (F to H) Representative immunostaining at week 14 showing (F) fibrotic area (Col1, yellow), bronchiolar epithelial cells (E-cadherin (E-cad), green) and AEC2s and bronchiolar epithelial cells (SP-C, red), (G) fibrotic area (Col1, white), PDGFRα(red) and PDGFRβ(green) and (H) Ki-67 (red), PDGFRα(green) and PDGFRβ (white). Scale bars = 50 μm. All specimens for immunostaining were obtained at week 14.

### Hyperplastic bronchiolar cells with invasive phenotype in the damaged lung

Hyperplastic epithelial cells were also prominent and widely distributed after BMS induction (Fig 5A). Co-staining of these E-cadherin-expressing cells for matrix metalloproteinase-7 (MMP7) and the N-terminal proteolytic fragments of laminin γ2 chain (γ2pf) was performed to assess how these cells might migrate or expand in the damaged lung (Miyazaki et al 2016). Although epithelial cells in the acute phase did not express these markers, MMP7 and γ2pf co-expression with E-cadherin was clearly evident after the intermediate phase (Figs 5A, 5B). This was associated with destruction of the basement membrane and cell migration into the alveolar space.

**Figure 5.**
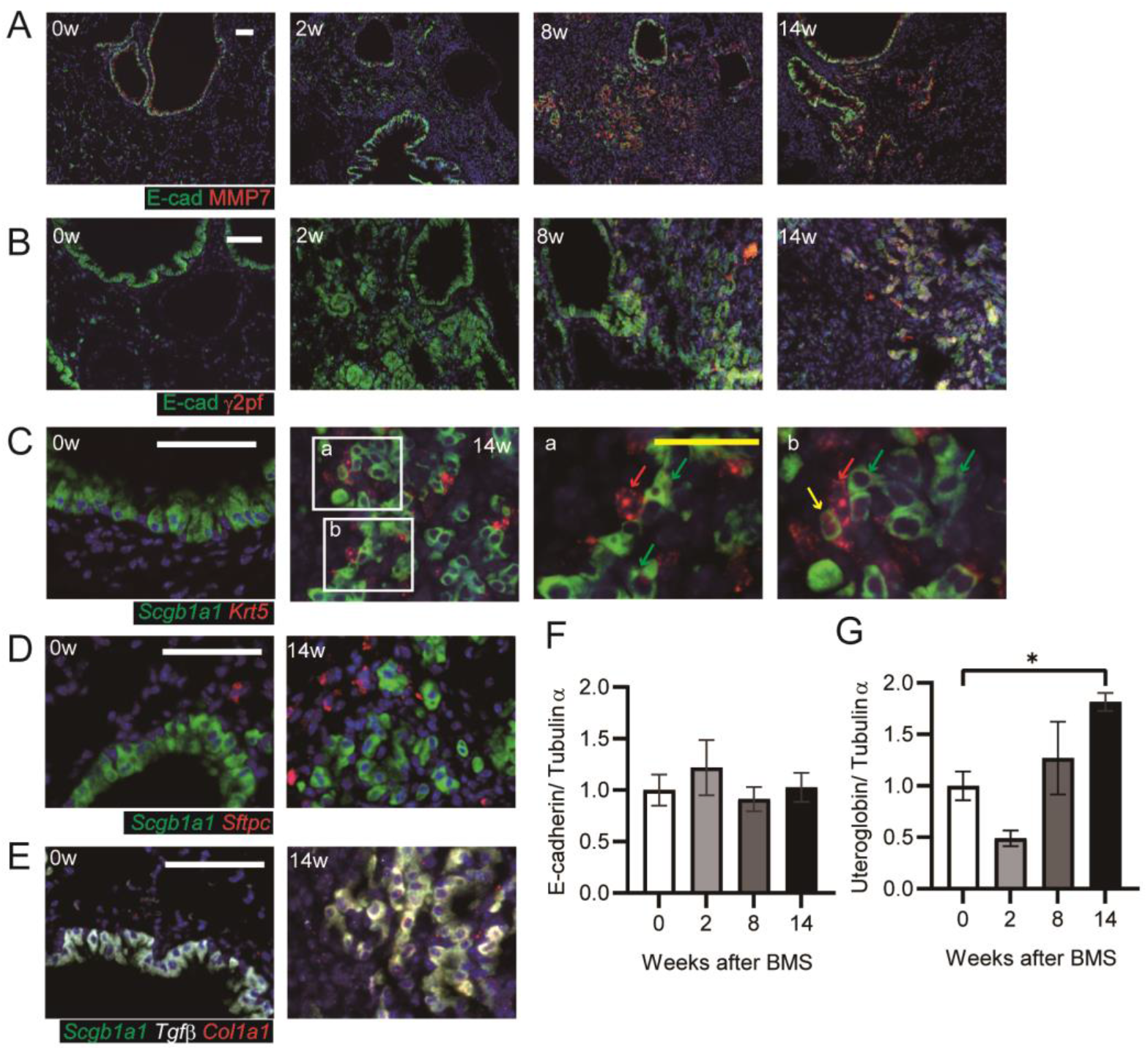
Invasive bronchiolar epithelial cells in chronic phase. Immunostaining for invasive bronchiolar epithelial cells using (A) E-cadherin (E-cad, green) and MMP7 (red) or (B) E-cadherin (E-cad, green) and N-terminal proteolytic fragments of Laminin, γ2pf (red). (C-E) *In situ* hybridization for invasive bronchiolar epithelial cells at weeks 0 and 14. (C) *Scgb1a1* (green) and *Krt5* (red), (D) *Scgb1a1*(green) and *Sftpc* (red), and (E) *Scgb1a1* (green), *Tgfβ1* (white), and *Col1a1* (red). Each arrow indicates *Krt5* (red), *Scgb1a1* (green), and *Krt5* and *Scgb1a1* (yellow) - positive cells in C a and b. Scale bars = 50 μm (white) and 25 μm (yellow). (F and G) E-cadherin (F) and uteroglobin (G) expression determined by WB and ImageJ, Fiji. Data are presented means ± S.E. of three mice. Asterisks show \**P* < 0.05, compared with week 0.

These invasive cells showed positive staining for secretoglobin, family 1A1 (*Scgb1a1*, which encodes uteroglobulin) and/or keratin 5 (*Krt5*), markers of club and basal cells, respectively (Fig 5C) (Hogan et al 2014). *Scgb1a1*-positive invasive cells also expressed fibrotic markers transforming growth factor-β1 (*Tgf-β1*) and collagen1A1 (*Col1a1*), suggesting the spread of fibrosis into areas of epithelial hyperplasia. These cells differed from bronchioalveolar stem cells (BASCs) because of this more fibrotic phenotype and less *Sftpc* expression (Figs 5D, 5E) (Jiang et al 2020, Kim et al 2005).

We next performed WB for total E-cadherin and uteroglobin expression in the lung. Uteroglobin expression, but not E-cadherin was increased at week 14 (Figs 5F, 5G). This suggested that the number of invasive epithelial cells was increased while the cell-to-cell contact of bronchiolar epithelial cells was not increased. Next, we tested whether these cells had undergone EMT by performing immunohistochemistry for the fibroblast marker S100A4. Evidence of E-cadherin and S100A4 co-staining was apparent at week 14 only (Fig S8). Taken together, these data suggest migratory epithelial cells lose their polarity, and become dysplastic with or without EMT.

### Poor reconstitution of blood vessels because of lack of CD31-positive endothelial cells

Poor vascularization results in pulmonary hypertension in patients with IPF (Nadrous et al 2005). We analyzed the failure of reconstitution of blood vessels after destruction of the lung structure following BMS administration. Initially, we investigated the localization of VEGF and PDGFRβ as key proteins related to vascularization. There was no appropriate blood vessel formation after BMS administration (Fig 6A). VEGF expression was evident in both normal as well as invasive bronchiolar epithelial cells (Figs 6A, 6B) (Fehrenbach et al 1999). In the fibrotic regions at week 14, PDGFRβ-positive fibroblasts aggregated in regions distinct from areas of VEGF-positive staining, while CD31 expression was low or almost undetectable (Fig 6A) (Renzoni et al 2003). Whole lung expression of CD31 and VEGF was also examined by WB, along with pigment epithelial derived growth factor (PEDF), which opposes angiogenesis. Acute BMS-induced decreases in VEGF and CD31 expression were not recovered by the intermediate or chronic phases (Figs 6C, 6D). A transient increase in PEDF expression at week 2 returned to near control levels at weeks 8 and 14 (Fig 6E). Further histopathological features were evident in the hyperplastic interstitial area. Decreased numbers of AEC1 and AEC2 were seen adjacent to hyperplastic epithelial cells (area a in Fig 6F); Blood vessels and endothelial cells remained in the interstitial area, but CD31 expression was low (area b in Fig 6F). Poor micro-vascular structure in the chronic phase was confirmed from micro-CT images of the lung vasculature filled ex vivo with Microfil. At 14 weeks post-BMS, there were fewer subpleural capillaries, and decreased volume of blood vessels (Figs 6G, 6H). Overall, the marked loss of endothelial cells in the lung may result in poor reconstitution of blood vessels, despite the aggregation of PDGFRβ-positive cells, which may have pericyte functions, in the fibrotic area.

**Figure 6.**
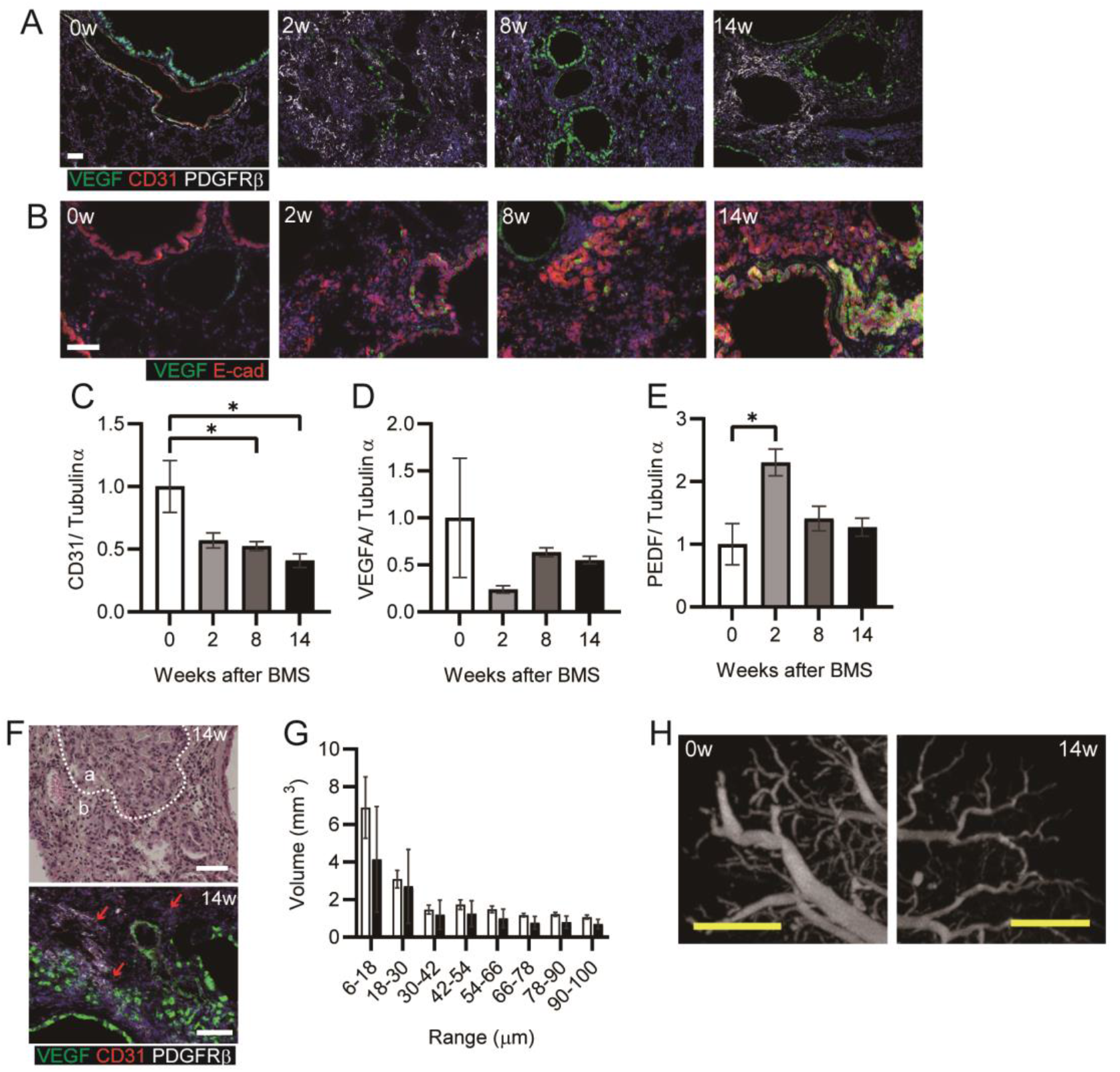
Poor reconstitution of blood vessels in chronic phase. (A) Immunohistochemical staining for VEGF (green), CD31 (red), and PDGFRβ (white) or (B) VEGF (green) and E-cadherin (Ecad, red) at weeks 0, 2, 8, and 14. (C) CD31, (D) VEGFA, and (E) PEDF expression was determined by WB and ImageJ, Fiji. Data are presented means ± S.E. of three mice for each week, 0, 2, 8 and 14. Asterisk shows \**P* < 0.05, compared with week 0. (F) Representative hematoxylin eosin staining (upper) and immunohistochemical staining for VEGF (green), CD31 (red), and PDGFRβ (white) (lower). “a” indicates hyperplastic area with bronchiolar epithelial cells, while “b” indicates hyperplastic interstitial area at week 14. Arrows indicate CD31-positive cells. (G) Representative subpleural images at weeks 0 and 14 showing blood vessels, visualized using Microfil and micro-CT. (H) Percentage blood vessel volume in the lung calculated after ex vivo Microfil application. Data are presented means ± S.E. of three mice for each week, 0 and 14. (A, B, F and G) Scale bars = 50 μm (white), 1 mm (yellow).

## DISCUSSION

A novel iUIP model using BMS as a stimulus was characterized by multi-step disease progression, most notably bimodal fibrosis with honeycombing. We propose that the application of this BMS method has increased the short half-life of bleomycin and/or provided a higher localized dose to induce these changes, but with lower mortality relative to previous studies using higher or repeated doses of intra-tracheal bleomycin (Satoh et al 2017, Tashiro et al 2017). IPF is a chronic, progressive interstitial lung disease that can occur as a result of prolonged IP. Severe cases of pneumonia and IPF can lead to death due to respiratory failure (Antoniou et al 2014). Fibrotic foci in IPF are often covered by a cuboidal lining epithelium, attributed to squamous metaplasia of epithelial cells (Batra et al 2018, Hashisako & Fukuoka 2015, Hogan et al 2014). In the current study, a similar metaplastic respiratory epithelium, which expressed E-cadherin, was observed after administration of BMS. The chronological features of multiple markers of disease progression in the iUIP mouse model are summarized in Fig 7. The clear pattern of bimodal fibrosis observed in the acute and chronic phases was similar to the histopathology of NSIP and UIP, respectively.

**Figure 7.**
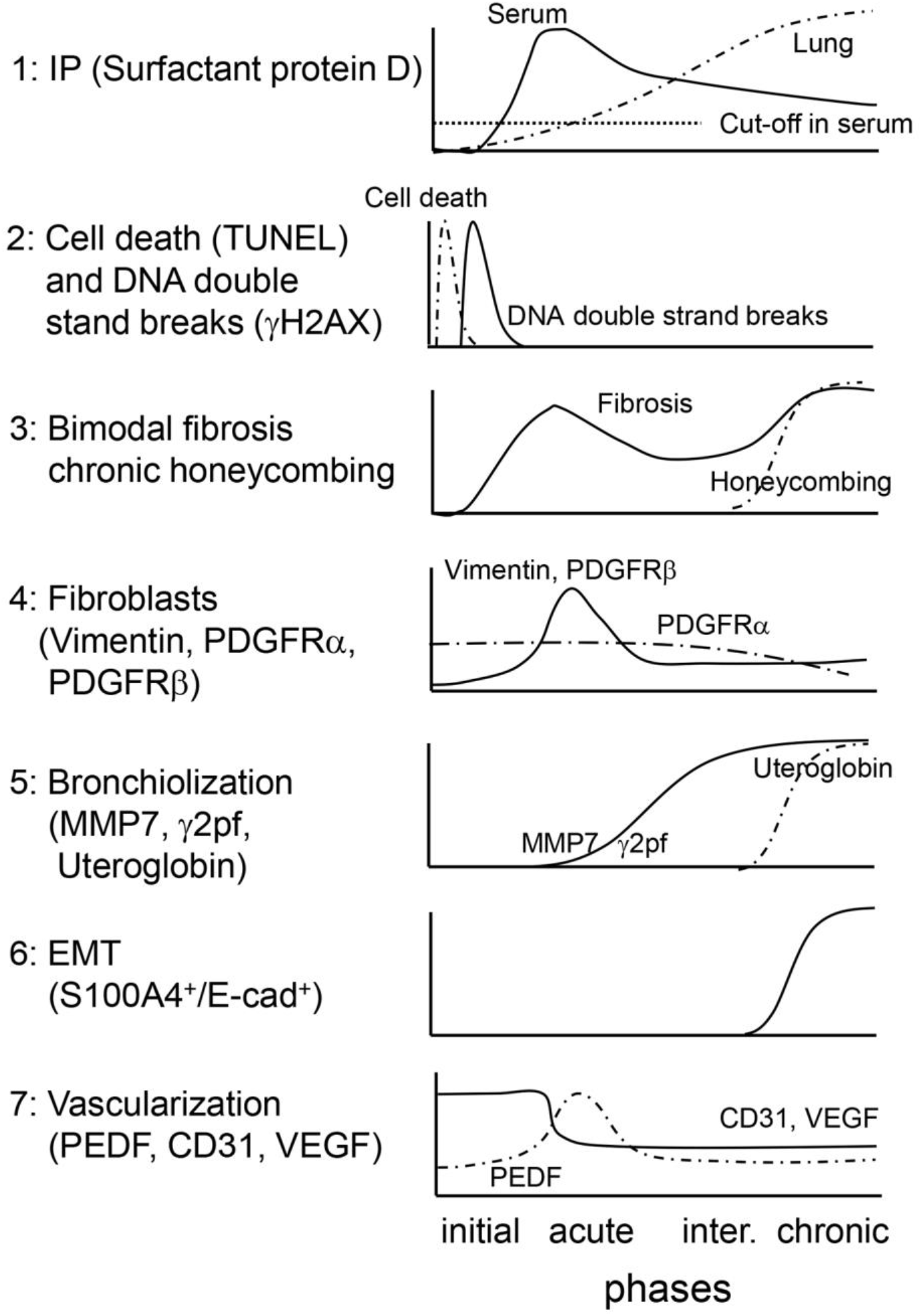
Chronological features in iUIP mouse model. 1) Serum SP-D levels immediately increased in acute phase, while total lung SP-D increased toward the chronic phase rather than the acute phase. 2) Cell death and DSBs in alveolar epithelial cells were induced by BMS administration. 3) Bimodal fibrosis was found in the acute and the chronic phases. Honeycombing was observed from the intermediate to chronic phases. 4) Fibroblast expansion rather than growth was evident in the acute phase. 5) Invasive epithelial cells, which expressed MMP7 and γ2pf, were observed in the intermediate and the chronic phases. 6) Some invasive epithelial cells expressed both E-cadherin and S100A4, suggesting that EMT occurring in the intermediate and chronic phases. 7) Poor vascularization was evidenced by reduction of CD31 and VEGF expression throughout the entire period after BMS administration. PEDF expression was upregulated only in the acute phase.

Bleomycin-induced DNA damage in the acute phase was limited to epithelial cells, with no overt DSBs detected in fibroblasts. This suggests that the secondary fibrosis with honeycombing may be indirectly regulated by these damaged epithelial cells in addition to fibroblasts. This would implicate NSIP as a previous stage of UIP, a possibility that could not be previously explored in conventional models, where chronic IP is not commonly established without repeated bleomycin dosing (Degryse et al 2010, Tashiro et al 2017).

During the acute phase, bronchiolar epithelial cells expressing a putative EMT biomarker MMP7 rather than the migratory marker S100A4 extended into the lung interstitium (Kropski et al 2015, Rosas et al 2008, White et al 2016). Enhanced migration from the area of hyperplastic bronchiolar epithelia was consistent with elevated MMP7 expression leading to destruction of the basement protein laminin. Expression of γ2pf, an additional invasive marker commonly seen in cancer cells, was increased in MMP7-positive cells in the intermediate and chronic phases. Thus, loss of cell-to-cell contact may cause hyperplastic alveoli to migrate into the lung interstitium to increase bronchiolization during the progression of secondary fibrosis. Of note, expression of MMP7 was elevated along with spontaneous lung fibrosis in a *Sftpc* mutant expressing mouse model (Nureki et al 2018).

*Krt5* expression was relatively weaker by week 14 in *Scgb1a1*-positive invasive cells, they may be similar to lineage-negative epithelial stem/ progenitor cells (LNEPs) or multipotent progenitor (Hogan et al 2014, Vaughan et al 2015). While this suggested that the cells had become dysplastic, they were different from squamous cell carcinoma in this stage. In a single cell analysis of epithelial cells in patients with IPF, the predominant cells were progenitor subpopulations, which express *Scgb1a1, Krt5, Krt8, and Trp63* (including ΔNp63)*, and TGF-b* as shown in the current model. The presence of these invasive epithelial cells may lead to failure to regenerate the lung epithelium during chronic fibrosis (Jiang et al 2020, Vaughan et al 2015). Further characterization of invasive epithelial cells in the intermediate and the chronic phases is required to implicate NSIP and DNA damage in epithelial cells as a requirement for the continuous pathological changes that contribute to chronic disease progression in some patients with IPF. Isolation of bronchiolar epithelia in specific areas of the lung using lineage tracing techniques will reveal patterns of gene expression to provide insights into potential factors that promote fibrosis.

Poor vascularization persisted from acute to chronic phases in BMS-induced D1CCxD1BC tg mice. The expression of pro-angiogenic VEGF was decreased at acute phase, while the corresponding inhibitor factor PEDF was transiently increased. It remains difficult to estimate the amount of active rather than total TGF-β, which is also a putative regulator for PEDF. Acute inflammation by bleomycin could affect the activation of TGF-β directly and/or indirectly through proteinase activation, resulting in fibrosis and vascular destruction. Although there was a persistent decrease in expression of CD31 simultaneously with VEGF during acute phase, this molecule may be more important for the transmigration of lymphocytes such as neutrophils, rather than for vascularization (Dangerfield et al 2002). These combined findings suggest a complex interplay between the epithelium, fibroblasts and endothelium in the initiation and progression of fibrosis. However, whether these observations are common phenomenon induced by bleomycin treatment or whether this poor vascularization persists only in BMS-induced D1CC×D1BC mice requires further investigation.

In summary, we have demonstrated that D1CC×D1BC mice develop acute IP, bimodal fibrosis, and chronic honeycombing in response to BMS. Critically, chronic IP and fibrosis were notably absent when intra-tracheal bleomycin alone was given to D1CC×D1BC mice or when BMS was administered to DBA/1J mice, demonstrating the importance of both the method of administration and the transgene in this model. It will be of interest to determine whether other bleomycin-susceptible mouse strains, such as the more commonly used C57BL/6 mice, have a similar amplification of the response to a single intra-tracheal dose of bleomycin when administered as BMS, as this could support the widespread use of this novel method of administration. While it remains clear that the mechanisms underlying fibrosis are complex, our iUIP model will facilitate further research on chronic IP and potential targets for new drug therapy.

## AUTHOR CONTRIBUTIONS

Conceptualization: S.K. Data curation: Y.M., S.K. Formal analysis: Y.M., M.L., J.B, S.K. Methodology: Y.M., M.L., J.B, S.K. Supervision: J.B, S.K. Visualization: Y.M., M.L., J.B, S.K. Writing-original draft: Y.M., S.K. Writing-review and editing: Y.M., M.L., J.B, S.K

## ACKNOWLEDMENTS

We thank for Drs. M. Murata and T. Numano for help in all aspects of this work. We acknowledge the assistance of Dr. Andre Tan from Histoindex for training in Genesis ^®^ 200 imaging and Fibroindex analysis.

## REFERENCES

Adamson IY. 1976. Pulmonary toxicity of bleomycin. Environmental health perspectives. 16:119–126. doi:10.1289/ehp.7616119

Antoniou KM, Margaritopoulos GA, Tomassetti S, Bonella F, Costabel U, Poletti V. 2014. Interstitial lung disease. Eur Respir Rev. 23(131):40–54. doi:10.1183/09059180.00009113

Ashcroft T, Simpson JM, Timbrell V. 1988. Simple method of estimating severity of pulmonary fibrosis on a numerical scale. J Clin Pathol. 41(4):467–470.

B. MB, Lawson WE, Oury TD, Sisson TH, Raghavendran K, Hogaboam CM. 2013. Animal models of fibrotic lung disease. American journal of respiratory cell and molecular biology. 49(2):167–179. doi:10.1165/rcmb.2013-0094TR

Batra K, Butt Y, Gokaslan T, Burguete D, Glazer C, Torrealba JR. 2018. Pathology and radiology correlation of idiopathic interstitial pneumonias. Hum Pathol. 72:1–17. doi:10.1016/j.humpath.2017.11.009

Borzone G, Moreno R, Urrea R, Meneses M, Oyarzun M, Lisboa C. 2001. Bleomycin-induced chronic lung damage does not resemble human idiopathic pulmonary fibrosis. American journal of respiratory and critical care medicine. 163(7):1648–1653. doi:10.1164/ajrccm.163.7.2006132

Cooper JA, Jr. 2000. Pulmonary fibrosis: Pathways are slowly coming into light. American journal of respiratory cell and molecular biology. 22(5):520–523. doi:10.1165/ajrcmb.22.5.f185

Dangerfield J, Larbi KY, Huang MT, Dewar A, Nourshargh S. 2002. Pecam-1 (cd31) homophilic interaction up-regulates alpha6beta1 on transmigrated neutrophils in vivo and plays a functional role in the ability of alpha6 integrins to mediate leukocyte migration through the perivascular basement membrane. The Journal of experimental medicine. 196(9):1201–1211. doi:10.1084/jem.20020324

Degryse AL, Tanjore H, Xu XC, Polosukhin VV, Jones BR, Boomershine CS, Ortiz C, Sherrill TP, McMahon FB, Gleaves LA, et al. 2011. Tgfbeta signaling in lung epithelium regulates bleomycin-induced alveolar injury and fibroblast recruitment. American journal of physiology Lung cellular and molecular physiology. 300(6):L887–897. doi:10.1152/ajplung.00397.2010

Degryse AL, Tanjore H, Xu XC, Polosukhin VV, Jones BR, McMahon FB, Gleaves LA, Blackwell TS, Lawson WE. 2010. Repetitive intratracheal bleomycin models several features of idiopathic pulmonary fibrosis. American journal of physiology Lung cellular and molecular physiology. 299(4):L442–452. doi:10.1152/ajplung.00026.2010

Dorr RT. 1992. Bleomycin pharmacology: Mechanism of action and resistance, and clinical pharmacokinetics. Semin Oncol. 19(2 Suppl 5):3–8.

Farkas L, Farkas D, Ask K, Moller A, Gauldie J, Margetts P, Inman M, Kolb M. 2009. Vegf ameliorates pulmonary hypertension through inhibition of endothelial apoptosis in experimental lung fibrosis in rats. The Journal of clinical investigation. 119(5):1298–1311. doi:10.1172/JCI36136

Fehrenbach H, Kasper M, Haase M, Schuh D, Muller M. 1999. Differential immunolocalization of vegf in rat and human adult lung, and in experimental rat lung fibrosis: Light, fluorescence, and electron microscopy. The Anatomical record. 254(1):61–73. doi:10.1002/(SICI)1097-0185(19990101)254:1<61::AID-AR8>3.0.CO;2-D

Flaherty KR, Wells AU, Cottin V, Devaraj A, Walsh SLF, Inoue Y, Richeldi L, Kolb M, Tetzlaff K, Stowasser S, et al. 2019. Nintedanib in progressive fibrosing interstitial lung diseases. The New England journal of medicine. 381(18):1718–1727. doi:10.1056/NEJMoa1908681

Fontes JD, Kanazawa S, Nekrep N, Peterlin BM. 1999. The class ii transactivator ciita is a transcriptional integrator. Microbes Infect. 1(11):863–869. doi:10.1016/s1286-4579(99)00232-4

Goh GB, Leow WQ, Liang S, Wan WK, Lim TKH, Tan CK, Chang PE. 2019. Quantification of hepatic steatosis in chronic liver disease using novel automated method of second harmonic generation and two-photon excited fluorescence. Scientific reports. 9(1):2975. doi:10.1038/s41598-019-39783-1

Hagimoto N, Kuwano K, Nomoto Y, Kunitake R, Hara N. 1997. Apoptosis and expression of fas/fas ligand mrna in bleomycin-induced pulmonary fibrosis in mice. American journal of respiratory cell and molecular biology. 16(1):91–101. doi:10.1165/ajrcmb.16.1.8998084

Hashisako M, Fukuoka J. 2015. Pathology of idiopathic interstitial pneumonias. Clin Med Insights Circ Respir Pulm Med. 9(Suppl 1):123–133. doi:10.4137/CCRPM.S23320

Hay J, Shahzeidi S, Laurent G. 1991. Mechanisms of bleomycin-induced lung damage. Archives of toxicology. 65(2):81–94.

Hogan BL, Barkauskas CE, Chapman HA, Epstein JA, Jain R, Hsia CC, Niklason L, Calle E, Le A, Randell SH, et al. 2014. Repair and regeneration of the respiratory system: Complexity, plasticity, and mechanisms of lung stem cell function. Cell stem cell. 15(2):123–138. doi:10.1016/j.stem.2014.07.012

Hubner RH, Gitter W, El Mokhtari NE, Mathiak M, Both M, Bolte H, Freitag-Wolf S, Bewig B. 2008. Standardized quantification of pulmonary fibrosis in histological samples. Biotechniques. 44(4):507–511, 514-507. doi:10.2144/000112729

Jenkins RG, Moore BB, Chambers RC, Eickelberg O, Konigshoff M, Kolb M, Laurent GJ, Nanthakumar CB, Olman MA, Pardo A, et al. 2017. An official american thoracic society workshop report: Use of animal models for the preclinical assessment of potential therapies for pulmonary fibrosis. American journal of respiratory cell and molecular biology. 56(5):667–679. doi:10.1165/rcmb.2017-0096ST

Jiang P, Gil de Rubio R, Hrycaj SM, Gurczynski SJ, Riemondy KA, Moore BB, Omary MB, Ridge KM, Zemans RL. 2020. Ineffectual type 2-to-type 1 alveolar epithelial cell differentiation in idiopathic pulmonary fibrosis: Persistence of the krt8(hi) transitional state. American journal of respiratory and critical care medicine. 201(11):1443–1447. doi:10.1164/rccm.201909-1726LE

Kanazawa S, Okamoto T, Peterlin BM. 2000. Tat competes with ciita for the binding to p-tefb and blocks the expression of mhc class ii genes in hiv infection. Immunity. 12(1):61–70. doi:10.1016/s1074-7613(00)80159-4

Kanazawa S, Ota S, Sekine C, Tada T, Otsuka T, Okamoto T, Sonderstrup G, Peterlin BM. 2006. Aberrant mhc class ii expression in mouse joints leads to arthritis with extraarticular manifestations similar to rheumatoid arthritis. Proceedings of the National Academy of Sciences of the United States of America. 103(39):14465–14470. doi:10.1073/pnas.0606450103

Kim CF, Jackson EL, Woolfenden AE, Lawrence S, Babar I, Vogel S, Crowley D, Bronson RT, Jacks T. 2005. Identification of bronchioalveolar stem cells in normal lung and lung cancer. Cell. 121(6):823–835. doi:10.1016/j.cell.2005.03.032

King TE, Jr., Bradford WZ, Castro-Bernardini S, Fagan EA, Glaspole I, Glassberg MK, Gorina E, Hopkins PM, Kardatzke D, Lancaster L, et al. 2014. A phase 3 trial of pirfenidone in patients with idiopathic pulmonary fibrosis. The New England journal of medicine. 370(22):2083–2092. doi:10.1056/NEJMoa1402582

Kropski JA, Pritchett JM, Zoz DF, Crossno PF, Markin C, Garnett ET, Degryse AL, Mitchell DB, Polosukhin VV, Rickman OB, et al. 2015. Extensive phenotyping of individuals at risk for familial interstitial pneumonia reveals clues to the pathogenesis of interstitial lung disease. American journal of respiratory and critical care medicine. 191(4):417–426. doi:10.1164/rccm.201406-1162OC

Limjunyawong N, Mitzner W, Horton MR. 2014. A mouse model of chronic idiopathic pulmonary fibrosis. Physiol Rep. 2(2):e00249. doi:10.1002/phy2.249

McElroy MC, Kasper M. 2004. The use of alveolar epithelial type i cell-selective markers to investigate lung injury and repair. The European respiratory journal: official journal of the European Society for Clinical Respiratory Physiology. 24(4):664–673. doi:10.1183/09031936.04.00096003

Miura Y, Kanazawa S. 2020. Osteochondrogenesis derived from synovial fibroblasts in inflammatory arthritis model. Inflamm Regen. 40:7. doi:10.1186/s41232-020-00115-w

Miura Y, Ota S, Peterlin M, McDevitt G, Kanazawa S. 2019. A subpopulation of synovial fibroblasts leads to osteochondrogenesis in a mouse model of chronic inflammatory rheumatoid arthritis. JBMR Plus. 3(6):e10132. doi:10.1002/jbm4.10132

Miyazaki K, Oyanagi J, Sugino A, Sato H, Yokose T, Nakayama H, Miyagi Y. 2016. Highly sensitive detection of invasive lung cancer cells by novel antibody against amino-terminal domain of laminin gamma2 chain. Cancer Sci. 107(12):1909–1918. doi:10.1111/cas.13089

Moeller A, Ask K, Warburton D, Gauldie J, Kolb M. 2008. The bleomycin animal model: A useful tool to investigate treatment options for idiopathic pulmonary fibrosis? The international journal of biochemistry & cell biology. 40(3):362–382. doi:10.1016/j.biocel.2007.08.011

Mouratis MA, Aidinis V. 2011. Modeling pulmonary fibrosis with bleomycin. Curr Opin Pulm Med. 17(5):355–361. doi:10.1097/MCP.0b013e328349ac2b

Murata M, Otsuka M, Mizuno H, Shiratori M, Miyazaki S, Nagae H, Kanazawa S, Hamaoki M, Kuroki Y, Takahashi H. 2010. Development of an enzyme-linked immunosorbent assay for measurement of rat pulmonary surfactant protein d using monoclonal antibodies. Experimental lung research. 36(8):463–468. doi:10.3109/01902141003746371

Nadrous HF, Pellikka PA, Krowka MJ, Swanson KL, Chaowalit N, Decker PA, Ryu JH. 2005. Pulmonary hypertension in patients with idiopathic pulmonary fibrosis. Chest. 128(4):2393–2399. doi:10.1378/chest.128.4.2393

Nathan SD, Noble PW, Tuder RM. 2007. Idiopathic pulmonary fibrosis and pulmonary hypertension: Connecting the dots. American journal of respiratory and critical care medicine. 175(9):875–880. doi:10.1164/rccm.200608-1153CC

Nureki SI, Tomer Y, Venosa A, Katzen J, Russo SJ, Jamil S, Barrett M, Nguyen V, Kopp M, Mulugeta S, et al. 2018. Expression of mutant sftpc in murine alveolar epithelia drives spontaneous lung fibrosis. The Journal of clinical investigation. 128(9):4008–4024. doi:10.1172/JCI99287

Okada K, Kudo N, Niwa K, Yamamoto K. 2005. A basic study on sonoporation with microbubbles exposed to pulsed ultrasound. J Med Ultrason (2001). 32(1):3–11. doi:10.1007/s10396-005-0031-5

Organ L, Bacci B, Koumoundouros E, Barcham G, Milne M, Kimpton W, Samuel C, Snibson K. 2015. Structural and functional correlations in a large animal model of bleomycin-induced pulmonary fibrosis. BMC Pulm Med. 15:81. doi:10.1186/s12890-015-0071-6

Phillips MR, Moore SM, Shah M, Lee C, Lee YZ, Faber JE, McLean SE. 2017. A method for evaluating the murine pulmonary vasculature using micro-computed tomography. J Surg Res. 207:115–122. doi:10.1016/j.jss.2016.08.074

Raghu G, Remy-Jardin M, Myers JL, Richeldi L, Ryerson CJ, Lederer DJ, Behr J, Cottin V, Danoff SK, Morell F, et al. 2018. Diagnosis of idiopathic pulmonary fibrosis. An official ats/ers/jrs/alat clinical practice guideline. American journal of respiratory and critical care medicine. 198(5):e44–e68. doi:10.1164/rccm.201807-1255ST

Redente EF, Black BP, Backos DS, Bahadur AN, Humphries SM, Lynch DA, Tuder RM, Zemans RL, Riches DW. 2021. Persistent, progressive pulmonary fibrosis and epithelial remodeling in mice. American journal of respiratory cell and molecular biology. doi:10.1165/rcmb.2020-0542MA

Redente EF, Jacobsen KM, Solomon JJ, Lara AR, Faubel S, Keith RC, Henson PM, Downey GP, Riches DW. 2011. Age and sex dimorphisms contribute to the severity of bleomycin-induced lung injury and fibrosis. American journal of physiology Lung cellular and molecular physiology. 301(4):L510–518. doi:10.1152/ajplung.00122.2011

Renzoni EA, Walsh DA, Salmon M, Wells AU, Sestini P, Nicholson AG, Veeraraghavan S, Bishop AE, Romanska HM, Pantelidis P, et al. 2003. Interstitial vascularity in fibrosing alveolitis. American journal of respiratory and critical care medicine. 167(3):438–443. doi:10.1164/rccm.200202-135OC

Rosas IO, Richards TJ, Konishi K, Zhang Y, Gibson K, Lokshin AE, Lindell KO, Cisneros J, Macdonald SD, Pardo A, et al. 2008. Mmp1 and mmp7 as potential peripheral blood biomarkers in idiopathic pulmonary fibrosis. PLoS Med. 5(4):e93. doi:10.1371/journal.pmed.0050093

Satoh T, Nakagawa K, Sugihara F, Kuwahara R, Ashihara M, Yamane F, Minowa Y, Fukushima K, Ebina I, Yoshioka Y, et al. 2017. Identification of an atypical monocyte and committed progenitor involved in fibrosis. Nature. 541(7635):96–101. doi:10.1038/nature20611

Schindelin J, Arganda-Carreras I, Frise E, Kaynig V, Longair M, Pietzsch T, Preibisch S, Rueden C, Saalfeld S, Schmid B, et al. 2012. Fiji: An open-source platform for biological-image analysis. Nat Methods. 9(7):676–682. doi:10.1038/nmeth.2019

Schurgers E, Mertens F, Vanoirbeek JA, Put S, Mitera T, De Langhe E, Billiau A, Hoet PH, Nemery B, Verbeken E, et al. 2012. Pulmonary inflammation in mice with collagen-induced arthritis is conditioned by complete freund’s adjuvant and regulated by endogenous ifn-gamma. European journal of immunology. 42(12):3223–3234. doi:10.1002/eji.201242573

Smith M, Dalurzo M, Panse P, Parish J, Leslie K. 2013. Usual interstitial pneumonia-pattern fibrosis in surgical lung biopsies. Clinical, radiological and histopathological clues to aetiology. J Clin Pathol. 66(10):896–903. doi:10.1136/jclinpath-2013-201442

Tashiro J, Rubio GA, Limper AH, Williams K, Elliot SJ, Ninou I, Aidinis V, Tzouvelekis A, Glassberg MK. 2017. Exploring animal models that resemble idiopathic pulmonary fibrosis. Front Med (Lausanne). 4:118. doi:10.3389/fmed.2017.00118

Terasaki Y, Terasaki M, Kanazawa S, Kokuho N, Urushiyama H, Kajimoto Y, Kunugi S, Maruyama M, Akimoto T, Miura Y, et al. 2019. Effect of h2 treatment in a mouse model of rheumatoid arthritis-associated interstitial lung disease. J Cell Mol Med. 23(10):7043–7053. doi:10.1111/jcmm.14603

Vaughan AE, Brumwell AN, Xi Y, Gotts JE, Brownfield DG, Treutlein B, Tan K, Tan V, Liu FC, Looney MR, et al. 2015. Lineage-negative progenitors mobilize to regenerate lung epithelium after major injury. Nature. 517(7536):621–625. doi:10.1038/nature14112

White ES, Xia M, Murray S, Dyal R, Flaherty CM, Flaherty KR, Moore BB, Cheng L, Doyle TJ, Villalba J, et al. 2016. Plasma surfactant protein-d, matrix metalloproteinase-7, and osteopontin index distinguishes idiopathic pulmonary fibrosis from other idiopathic interstitial pneumonias. American journal of respiratory and critical care medicine. 194(10):1242–1251. doi:10.1164/rccm.201505-0862OC

Xenariou S, Liang HD, Griesenbach U, Zhu J, Farley R, Somerton L, Singh C, Jeffery PK, Scheule RK, Cheng SH, et al. 2010. Low-frequency ultrasound increases non-viral gene transfer to the mouse lung. Acta Biochim Biophys Sin (Shanghai). 42(1):45–51.

Yamaguchi H, Ishida Y, Hosomichi J, Suzuki JI, Usumi-Fujita R, Shimizu Y, Kaneko S, Ono T. 2017. A new approach to transfect nf-kappab decoy oligodeoxynucleotides into the periodontal tissue using the ultrasound-microbubble method. Int J Oral Sci. 9(2):80–86. doi:10.1038/ijos.2017.10

