## supplementary materials for "Bimodal fibrosis in a novel mouse model of bleomycin-induced usual interstitial pneumonia"

Authors and affiliations

Yoko Miura[1], Maggie Lam[2], Jane E. Bourke[2], and Satoshi Kanazawa[1]

[1] Department of Neurodevelopmental Disorder Genetics, Nagoya City University Graduate School of Medical Sciences, Nagoya, Aichi, Japan

[2]Department of Pharmacology, Biomedicine Discovery Institute, Monash University, Clayton, Australia

**Figure S1 Absence of IP with sonoporation and/or microbubbles without bleomycin, or with lower doses of BMS**

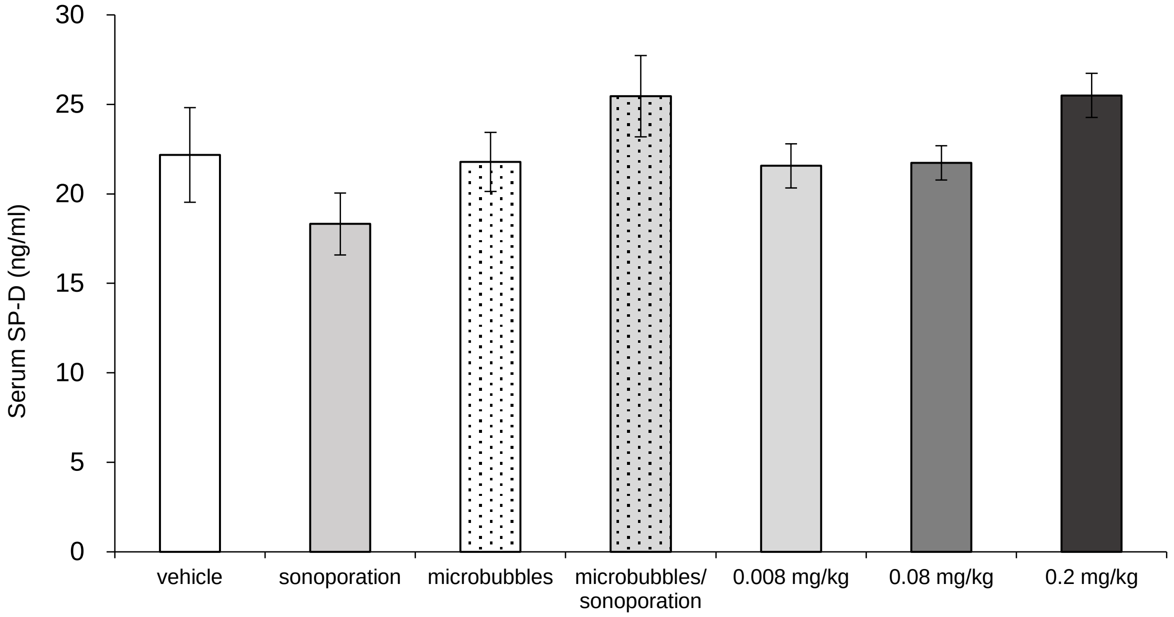


Sonoporation or microbubbles, alone and in combination, or lower doses of BMS (bleomycin 0.008, 0.8, 0.2 mg/kg) are not sufficient to induce IP in D1CC×D1BC mouse. Serum SP-D was measured by ELISA in samples collected at week 2 after BMS administration, corresponding to peak levels with BMS (1.28 mg/kg) in Figure 1. The results represent means ± S.E. of eight mice per group.

**Figure S2 TUNEL assay with conventional bleomycin method**

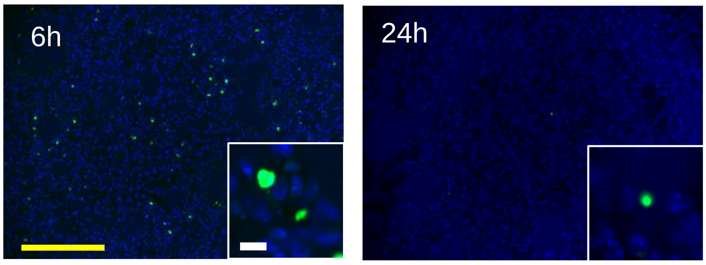


Intratracheal instillation of bleomycin (40µl/ mouse, 3.84 mg/kg, three times higher than in BMS method) was performed in D1CC×D1BC mice. Mice were sacrificed at 6 or 24h after bleomycin administration. Scale bars = 100 µm (yellow) and 10 µm (white).

**Figure S3 Infiltration of lymphoid cells by day 7 post-BMS, without systemic tissue damage.**

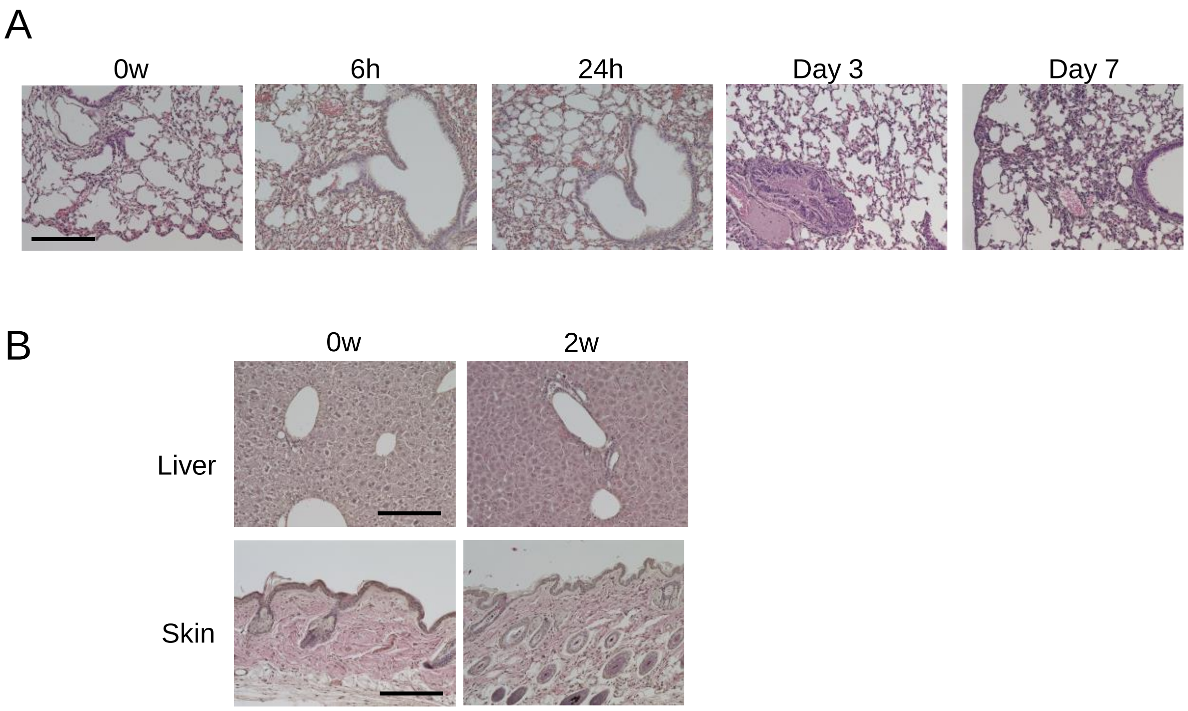


Histochemical staining with hematoxylin and eosin after BMS administration. (A) Lung sections at 0, 6, and 24 hours, and 3 and 7 days (B) Liver and skin sections from control mice (left) and at week 2 after BMS (right). There were no overt phenotypic changes in the liver or skin following bleomycin treatment. Scale bars = 100 μm.

**Figure S4 Honeycomb structure in the chronic phase of BMS model**

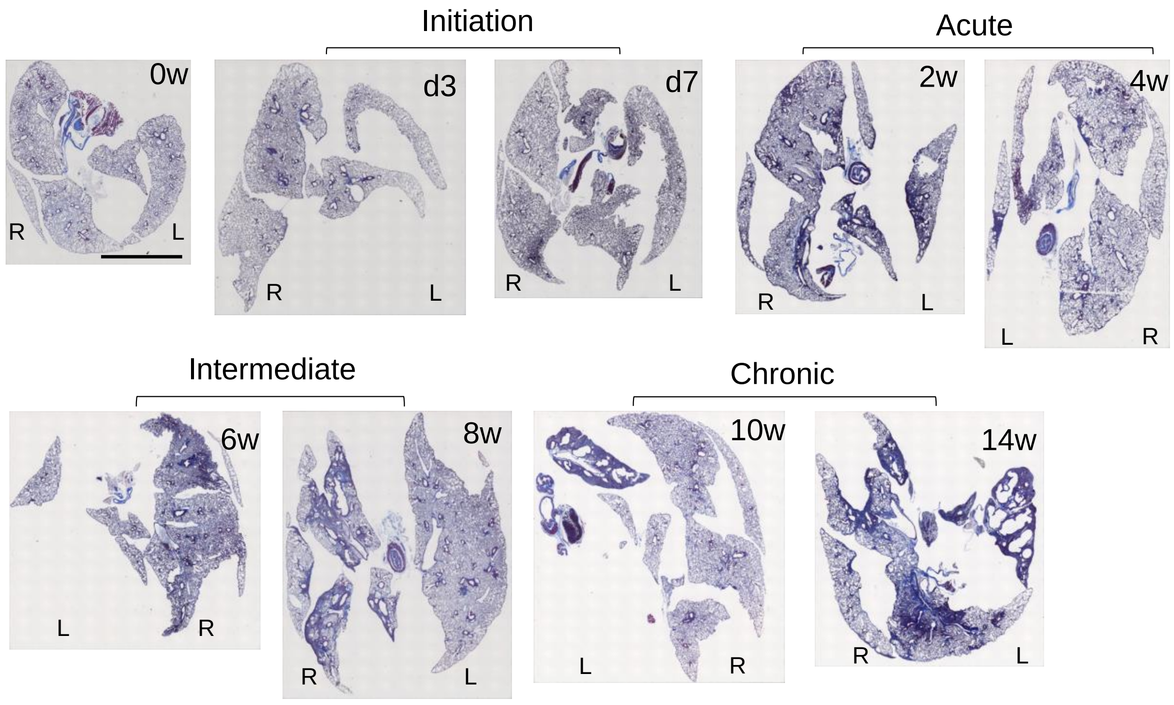


Representative macrographs of Masson’s trichrome staining at days 0, 3, 7 and weeks 2, 4, 6, 8, and 14 after BMS administration in whole lung sections from D1CC×D1BC mice. L: Left lobe, R: Right lobe. Scale bar = 5 mm.

**Figure S5 Severe inflammation in acute but not chronic phases of BMS model**

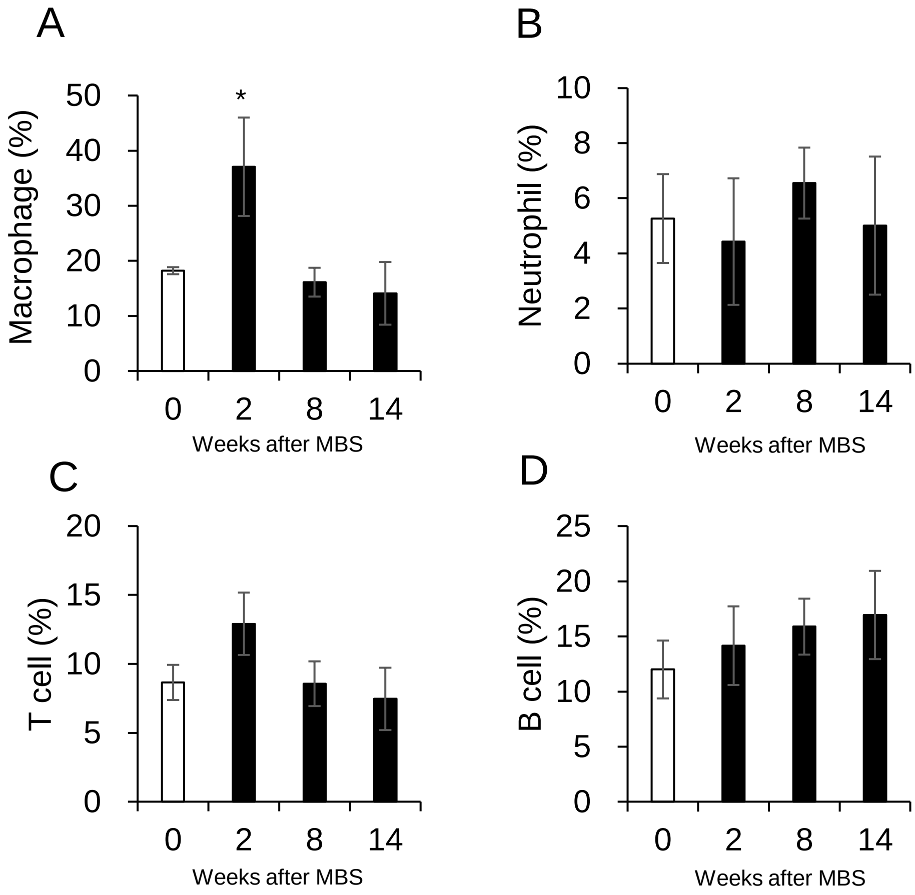


Percentages of lymphoid cells in whole lung images, determined by immunohistochemical staining for (A) Macrophage (F4/80), (B) Neutrophil (PAD4), (C) T cell (CD3), and (D) B cell (CD45R) at weeks 0, 2, 8, and 14 and calculated by ImageJ, Fiji. All results are represented as the means ± S.E. from lung sections from three mice for each timepoint. Asterisk shows $P < 0.05$, compared with 0 weeks.

**Figure S6 BMS induces mild chronic fibrosis in D1CC, D1BC, and DBA/1J mice.**

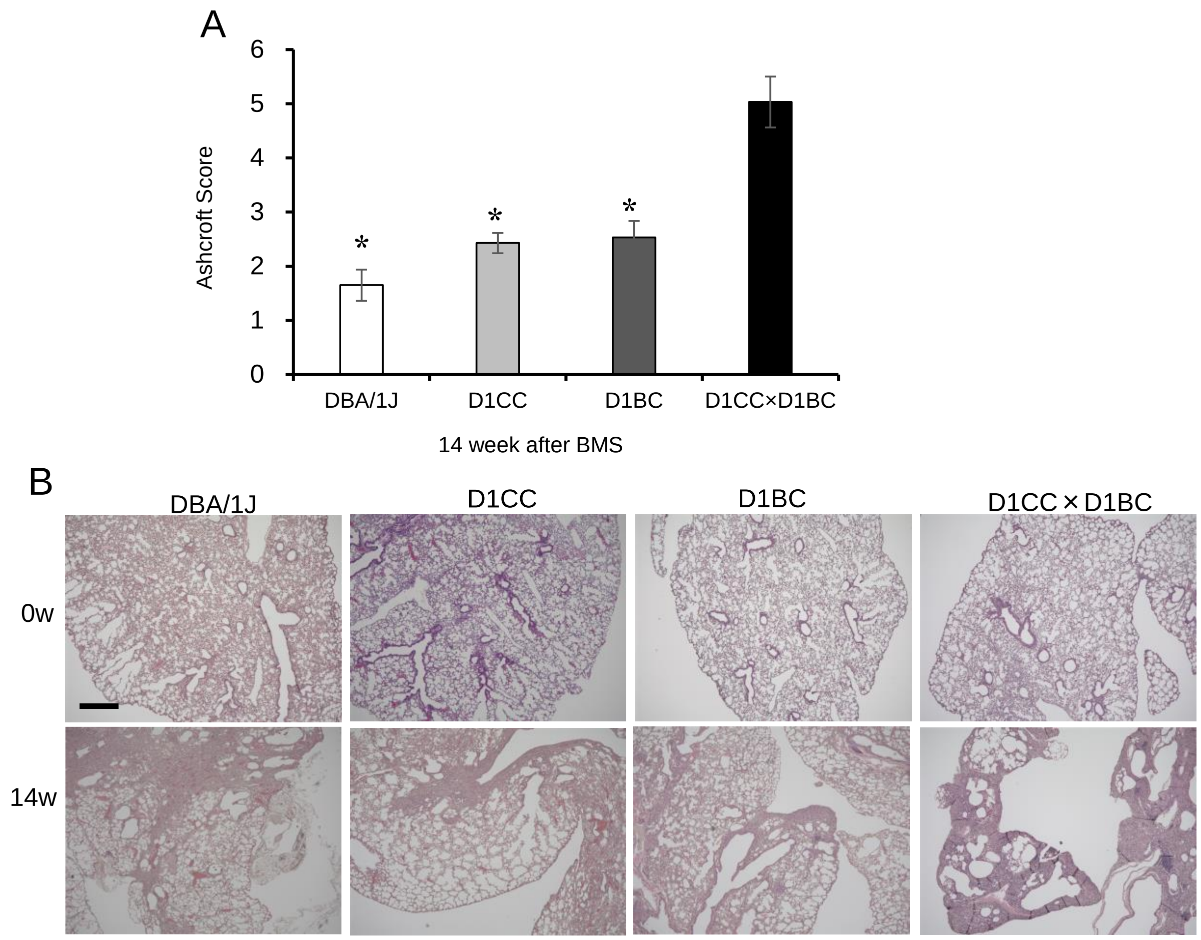


Weak IP progression in D1CC, D1BC alone, and their genetic background DBA/1J mice treated with BMS. (A) Quantitative analysis of Ashcroft score at week 14 for DBA/1J, D1CC, D1BC, and D1CC×D1BC mice (averaged from scores by two individuals, YM and SK, performed in a blinded manner). The results represent means ± S.E. from five mice (DBA/1J and D1CC) or three mice (D1BC and D1CC×D1BC) per group. Asterisk shows $P < 0.05$, compared with D1CC×D1BC mice. (B) Histochemical staining with hematoxylin and eosin. Scale bar = 500 μm.

**Figure S7 Elevated αSMA expression after BMS administration.**

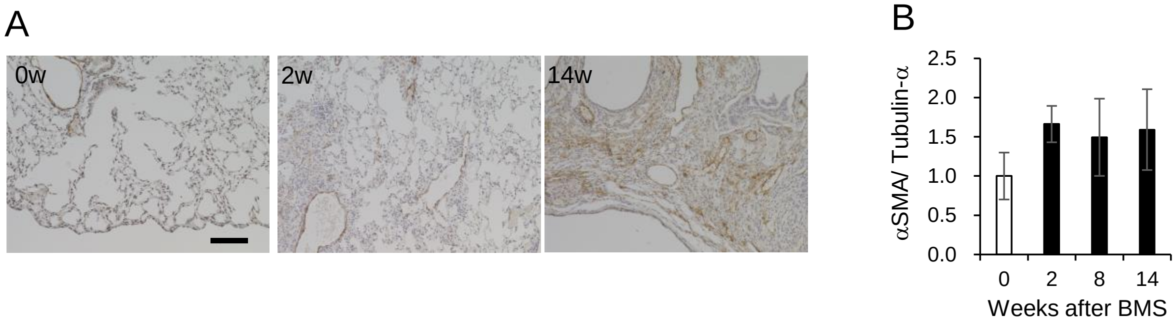


(A) Immunohistochemical staining for αSMA (DAB staining) at weeks 0, 2, and 14. Scale bar = 100 μm. (B) Expression of αSMA at weeks 0, 2, 8, and 14 was determined by WB and ImageJ, Fiji. Data are presented means ± S.E. of three mice.

**Figure S8 Invasive bronchiolar epithelial cells with less EMT in chronic phase of BMS model**

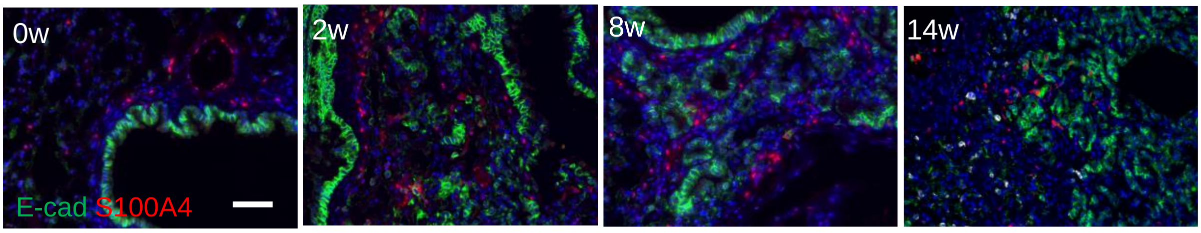


Immunostaining for epithelial cells using E-cadherin (green) and S100A4 (red) as an EMT marker (red). Scale bar = 50 μm.
